# OrfViralScan 3.0: An intuitive tool for the identification and tracking of open reading frames in viral genomes

**DOI:** 10.1101/2025.04.26.650794

**Authors:** Roberto Reinosa Fernández

## Abstract

The identification and analysis of open reading frames (ORFs) are fundamental steps in genome annotation, which requires accurate bioinformatics tools. OrfViralScan 3.0 is presented, a desktop application developed in Java (requires Java 11 to run), featuring an intuitive graphical user interface (GUI) designed to facilitate the annotation and tracking of ORFs.

The program offers functionalities to search for ORFs (initiated by ATG) in individual sequences (ORF Search), track specific ORFs based on length and location across multiple genomes or sequence sets (Track Specific ORF), preprocess FASTA files to standardize formatting (Preprocess File), and split large genomes into manageable fragments (Divide Into Fragments). The utility of OrfViralScan 3.0 is demonstrated through the analysis of the SARS-CoV-2 reference genome (NC_045512.2), the successful tracking of the Spike protein in 983 out of 1000 complete viral genomes, and the preparation of the Escherichia coli genome (NC_000913.3) for fragmented analysis. The software’s capabilities, limitations, and potential future applications are discussed. An example is also included featuring the SARS-CoV-2 Spike protein, showing the folded ORF obtained with OrfViralScan 3.0 using AlphaFold 3. The program’s source code is available on GitHub under the GNU GPLv3 license.

## 1. Introduction

Open reading frames (ORFs) represent sequences of genetic material with the potential to be translated into proteins [1]. In this context, ORFs are defined as beginning with a start codon (usually ATG, which encodes the amino acid methionine) and ending with a stop codon (TAA, TAG, or TGA) [1]. Accurate identification of ORFs is crucial for understanding an organism’s biology, predicting gene functions, and studying evolution [1]. OrfViralScan 3.0 focuses on viruses, whose genomes are often compact [2]. OrfViralScan 3.0 searches for ORFs starting with ATG and ending with one of the stop codons, without considering overlaps. This approach is generally sufficient to identify most ORFs in viral genomes. In eukaryotic genomes, more sophisticated methods are usually required due to the presence of introns.

Bioinformatics offers various tools for ORF prediction [3, 4], but often a combination of programs or custom scripts is needed for specific tasks, such as tracking a particular ORF across multiple viral sequences or performing general ORF searches in a sequence. To address these needs, OrfViralScan 3.0 has been developed as a Java-based program that provides an integrated and user-friendly solution. Its graphical interface simplifies the process for researchers without advanced programming experience.

**Figure 1.**
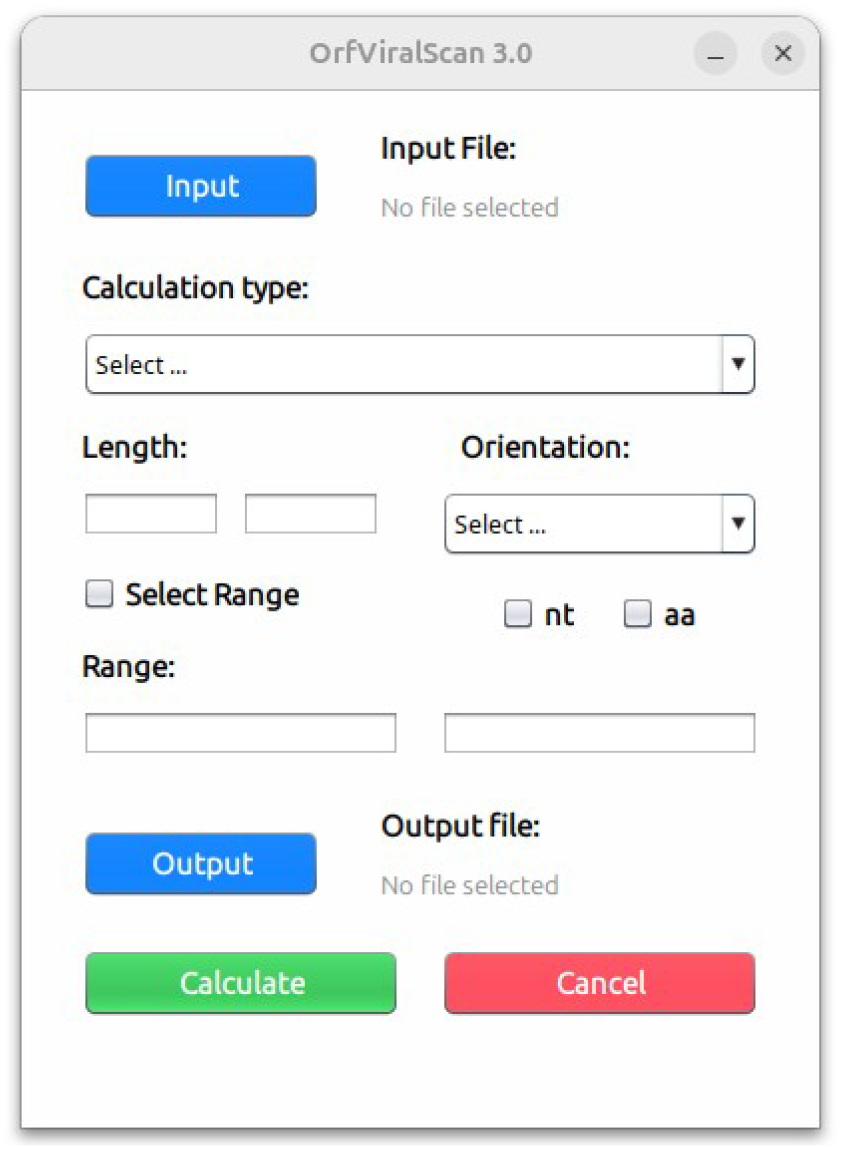
Graphical interface of OrfViralScan 3.0.

This article describes the functionalities of OrfViralScan 3.0 and presents case studies that illustrate its practical application in viral (SARS-CoV-2) and bacterial (E. coli) genomes. Although the program is not designed for bacterial genomes or very large genomes, it can still be used with them. This comes with certain limitations, such as increased computation time and the lack of Shine-Dalgarno region identification [5], which is very useful for predicting ORFs in bacterial genomes.

The source code of OrfViralScan 3.0, as well as the executable (JAR file), can be found at the following GitHub address: (https://github.com/roberto117343/OrfViralScan/). It is worth mentioning that the fact that OrfViralScan 3.0 is labeled as its third version is due to its foundation on two previous versions; however, significant changes have been made that justify the present work.

## 2. Program Description and Features

OrfViralScan 3.0 is a desktop application developed in Java using the OpenJDK. It requires Java version 11 to run. The program was developed with the Apache NetBeans 14 IDE on the Ubuntu 24.04.2 LTS operating system. It features a graphical user interface (GUI) designed to be intuitive and to provide easy access to its four main functionalities.

### 2.1. ORF Search

This functionality identifies non-overlapping ORFs from the first start codon (ATG) to the next stop codon (TAA, TAG, TGA) within a nucleotide sequence in a FASTA file [6].

**Input:** A FASTA file (preferably preprocessed) containing a single sequence.

**Parameters:** The user can choose to search for ORFs on the input strand, the antiparallel strand, or both. This must be selected in the “Orientation” section.

**Process:** The program scans the three reading frames for each selected strand. It identifies ORFs by finding the first start codon (ATG) and extending until the first stop codon (TAA, TAG, TGA) found in the same frame.

**Important:** Overlapping ORFs are not considered; if a start codon is found within an existing ORF in the same frame, a new ORF is not initiated from that point. The coordinates of ORFs found on the antiparallel strand correspond to their location on that strand.

**Output:** A text file (.txt) detailing the ORFs found. This file is divided into two sections: the first for nucleotide sequences and the second for amino acid sequences. For each ORF, the following information is provided:

- Reading frame (Frame 1, 2, or 3 and antiparallel frames).
- Unique identification number (#ID).
- Nucleotide sequence of the ORF (Sequence).
- Start and End positions in the nucleotide sequence.
- Length of the ORF in nucleotides (Length).

In the amino acid section, the program includes:

- Reading frame (Frame 1, 2, or 3 and antiparallel frames).
- Unique identification number (#ID), matching the nucleotide #ID for the same ORF.
- Amino acid sequence (Sequence).
- Length in amino acids (Length).
- Theoretical molecular mass in kilodaltons (Molecular Mass ∼ X kDa).

If the original nucleotide sequence contained ambiguous characters (‘N’) or any character other than A, C, T, G, resulting in a ‘?’ in the amino acid sequence, the molecular mass is indicated as -1, signaling that accurate calculation was not possible.

### 2.2. Track Specific ORF

Designed to locate a specific ORF of interest across multiple genomes, useful for phylogenetic or genetic variation studies.

**Input:** A preprocessed FASTA file, using “Preprocess File” as explained later, containing one or more sequences. An example would be complete SARS-CoV-2 genomes.

**Parameters:** A series of parameters that the user must specify:

- Length: Range of sequence length to search for the target ORF. This parameter can refer to nucleotides or amino acids depending on the search type.
- Orientation: Selects where to perform the search — Input Strand (sequences from the FASTA file), Antiparallel Strand (antiparallel to the input sequences), or Both (both strands).
- Sequence type: Check “nt” for nucleotides or “aa” for amino acids.
- Search Range (Select Range): This parameter is optional but recommended. It corresponds to the minimum and maximum position in the genome where an ORF is expected to be found.

This information can be obtained beforehand using “ORF Search” on a reference sequence.

**Process:** For each sequence in the input file, the program searches for an ORF that meets the specified parameter criteria. It uses the same detection logic (ATG to stop codon) as “ORF Search”. If a matching ORF is found, it is saved to the output file.

**Output:** A FASTA file containing the ORFs found. Each sequence in the output file corresponds to the ORF identified in one of the input sequences. The FASTA header of each recovered ORF includes the header of the original sequence it was derived from, and if the search was performed in amino acids, the calculated molecular mass (in kDa) is added to the header.

It is important to note that the ORF found for each sequence may not be correct if the search parameters are too lax or if there are two ORFs of similar length and position within the genome. It is recommended to edit the obtained file using programs like MEGA [7] to remove incorrectly identified ORFs. If too many incorrect ORFs are found, it is advisable to adjust the search parameters.

### 2.3. Preprocess File

This functionality is essential to ensure compatibility with the other features.

**Input:** A FASTA format file, which may contain one or multiple sequences.

**Process:** It reads the input FASTA file and converts it into a format where each sequence occupies exactly two lines: one line for the header (starting with ‘>‘) and the following line containing the entire sequence without internal line breaks.

**Output:** A FASTA file with the desired format.

This format is required for the “Track Specific ORF” and “Divide Into Fragments” functionalities, and recommended for “ORF Search,” as it prevents reading errors due to line breaks or empty lines.

### 2.4. Divide Into Fragments

A functionality that splits a very large sequence into different files of 100,000 nucleotides each. This helps ensure that “ORF Search” does not become too slow and potentially avoids memory limit issues.

**Input:** A preprocessed FASTA file with a single sequence (typically a very large genomic sequence).

**Process:** The input sequence is divided into non-overlapping fragments of 100,000 nucleotides. The last fragment may be shorter if the total length of the sequence is not an exact multiple of 100,000, which is usually the case.

**Output:** Multiple FASTA files, each containing a single fragment. The files are named sequentially (fragment_1.fasta, fragment_2.fasta, etc.). The user must specify the output folder where these files should be saved.

While OrfViralScan 3.0 is currently not designed to analyze genomes larger than average viral genomes, this functionality allows “ORF Search” to analyze each fragment individually. However, there are still limitations when predicting ORFs in genomes such as bacterial genomes, since Shine-Dalgarno regions or alternative start codons other than ATG are not considered.

## 3. Case Studies

### 3.1. Identification of ORFs in the Reference Genome of SARS-CoV-2

**Process:** The reference genome of SARS-CoV-2 (Wuhan-Hu-1), accession number NC_045512.2 [8], was downloaded in FASTA format from NCBI [9]. The file was processed using “Preprocess File” to ensure compatibility with other functionalities of the program. Afterwards, “ORF Search” was used, selecting the search only on the input strand.

**Results:** The program generated a TXT file listing the ORFs found. The output format is clear and provides the frame, ID, sequence, coordinates, and length for each nucleotide ORF, followed by the frame, ID, sequence, length, and molecular mass information for each amino acid sequence.

#### Output Analysis

Multiple ORFs were identified in the three reading frames of the input strand. Some of them are detailed below:

- **ID #2** (Frame 1, Start = 13768, End = 21555, Length = 7788 nt) is a very long ORF that does not directly correspond to the canonical ORF1ab (which starts much earlier, at position 266). This ORF found by the program starts at an internal methionine (position 13768) within ORF1ab [10] and extends to its stop codon (position 21555). This illustrates the limitation of strictly searching from the first ATG to the next stop codon without considering the overall genomic context or nested/alternative ORFs within polyproteins. This occurs due to a frameshift event.
- **ID #4** (Frame 1, Start = 25393, End = 26220, Length = 828 nt) corresponds to the ORF3a gene [10]. The calculated molecular mass for this protein is approximately 31.12 kDa.
- **ID #5** (Frame 1, Start = 26245, End = 26472, Length = 228 nt) corresponds to the Envelope (E) protein [10]. Molecular mass ∼8.36 kDa.
- **ID #7** (Frame 1, Start = 27394, End = 27759, Length = 366 nt) corresponds to the ORF7a [10]. Molecular mass ∼13.74 kDa.
- **ID #9** (Frame 2, Start = 266, End = 13483, Length = 13218 nt) represents the ORF1a polyprotein [10]. Molecular mass ∼ 489.99 kDa.
- **ID #20** (Frame 2, Start = 21536, End = 25384, Length = 3849 nt) corresponds to the Spike (S) protein [10]. Its calculated molecular mass is approximately 142.27 kDa. It should be noted that the predicted protein is 9 amino acids longer than the actual Spike protein.
- **ID #25** (Frame 2, Start = 28274, End = 29533, Length = 1260 nt) corresponds to the Nucleocapsid (N) protein [10]. Molecular mass ∼ 45.63 kDa.
- **ID #56** (Frame 3, Start = 26523, End = 27191, Length = 669 nt) corresponds to the Membrane (M) protein [10]. Molecular mass ∼ 25.15 kDa.

The ORFs presented here are just a small sample of those found. The analysis also identified shorter ORFs that may be artifacts or small peptides. The larger an ORF, the more likely it is to be translated into a functional protein. ORFs with fewer than 30 amino acids are filtered out and therefore not shown by the program.

### 3.2 Tracking the Spike Protein in 1000 SARS-CoV-2 Genomes

#### Process

Using the previously obtained information from “ORF Search” regarding the protein with ID #20 (Spike) shown in the previous step, the tracking was extended to a larger dataset. A total of 1000 complete SARS-CoV-2 genome sequences were downloaded from the NCBI Virus database [11]. The FASTA file was first processed with “Preprocess File”. Then, “Track Specific ORF” was used with the following parameters:

- **Length:** 1262–1302 amino acids (allowing slight variation around the 1282 amino acids of the ORF with ID #20, to capture natural variants).
- **Orientation:** Input Strand.
- **Sequence Type:** Amino acids (aa).
- **Range:** 19536–27384 nt (a broad range based on the previously obtained location, to ensure capture despite possible insertions/deletions in the genomes).

## Results

The program processed the 1000 genomes and generated an output FASTA file containing 983 sequences corresponding to the identified Spike protein ORF. Each header included the molecular mass in kDa.

The successful retrieval of 983/1000 (98.3%) Spike sequences demonstrates the effectiveness of “Track Specific ORF” for large-scale studies. Allowing slight variation in length (1262–1302 aa) was crucial for capturing natural variants, including those with a few additional amino acids at the start or end compared to the reference, as observed.

However, it is important to highlight a limitation: if a search range is not specified (or if it is too broad) and there are other ORFs nearby with similar lengths, the tool may retrieve incorrect ORFs. In such cases, manual verification is necessary, for example using MEGA software. The preprocessing functionality was essential to correctly load the input FASTA file.

**Figure 2.**
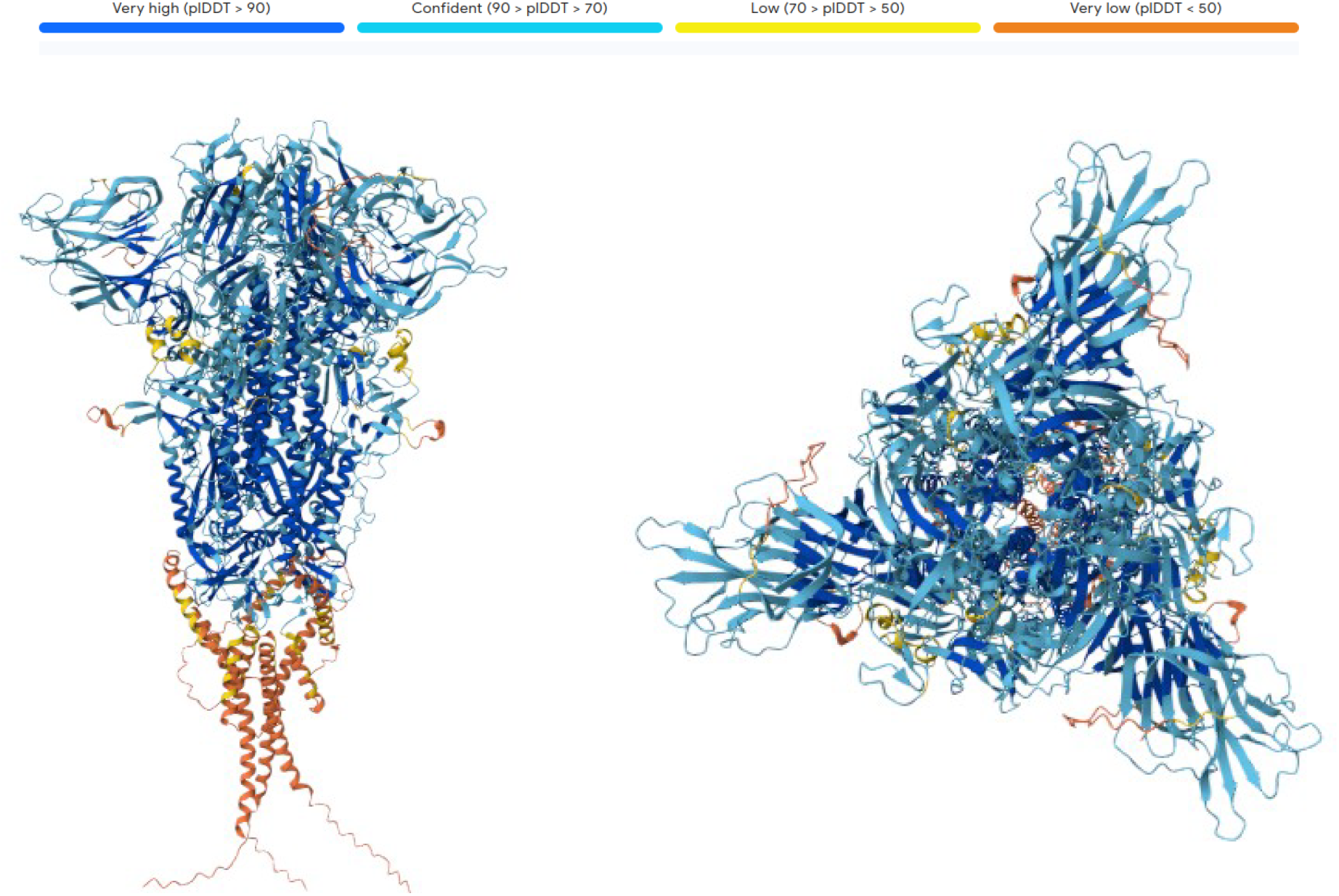
Three-dimensional structure predicted with AlphaFold 3 [12] using the sequence obtained from “ORF Search” (ID #20) corresponding to the Spike protein of SARS-CoV-2.

### 3.3 Preparation and Partial Analysis of the E. coli Genome

#### Process

The reference genome of Escherichia coli K-12 substr. MG1655 (NC_000913.3) [13], approximately 4.6 Mbp in size, was downloaded. Given its length, “Preprocess File” was first applied, followed by “Divide Into Fragments”. This generated 47 files, each containing genome fragments of 100,000 nucleotides, except for the last one, which is smaller. As a demonstration, the first fragment (fragment_1.fasta) was selected and analyzed using “ORF Search”, scanning for ORFs on both strands (Input Strand and Antiparallel Strand).

#### Results and Discussion

Running “ORF Search” on a 100,000 nt fragment was significantly faster than attempting to analyze the entire genome at once, validating the usefulness of the “Divide Into Fragments” functionality for large genomes. However, it is confirmed that “ORF Search”, even when applied to fragments, can be computationally intensive, especially when scanning both strands.

The output for fragment 1 showed ORFs on both strands and across all three reading frames, as expected for a bacterial genome. This fragment-based approach allows for manageable analysis but requires subsequent compilation of results for a global view of the genome.

The limitation of only recognizing ATG as a start codon remains. It must also be considered that fragmenting the genome can split an ORF, leading to incorrect predictions, and may shift the reading frame because each resulting fragment starts reading from nucleotide one in frame 1.

## 4. Discussion

OrfViralScan 3.0 emerges as a valuable and accessible tool for the scientific community, especially for those working with viral genomes. Its graphical user interface and logical workflow (Preprocess → Divide (optional) → Search → Track) simplify tasks that would otherwise require multiple tools or custom scripts.

### Advantages

- **Intuitive interface:** Facilitates use by researchers with varying levels of bioinformatics experience.
- **Integrated functionalities:** Combines ORF search and tracking within a single application.
- **Standard outputs:** “Track Specific ORF” generates output in FASTA format, ideal for programs like EpiMolbio [14], where comprehensive genetic variability analyses can be performed.
- **Portability:** Written in Java, it is potentially executable on any operating system with a compatible JVM (version 11).

## Limitations

- **Fixed Start Codon:** Only recognizes ATG as the start codon, ignoring alternative start codons that may be relevant.
- **Speed on Large Genomes:** “ORF Search” can be slow on very long sequences, although the “Divide Into Fragments” feature partially mitigates this issue.
- **Ambiguity in “Track Specific ORF”:** Without a well-defined search range and length constraints, it may retrieve incorrect ORFs if multiple ORFs meet the same criteria in a region. External validation is required.
- **Dependency on Java 11:** Requires a specific version of Java installed to ensure correct operation.

### Future Applications and Perspectives

The use of OrfViralScan 3.0 is planned for research in various fields of microbiology. Future versions of the software are expected to include options to search for alternative start codons, implement more sophisticated algorithms to detect ORFs and optimize search speed, and add alternative output formats for “ORF Search” to present information more canonically, as well as improvements in annotation.

## 5. Conclusion

OrfViralScan 3.0 is a robust and user-friendly tool that addresses specific needs in ORF analysis, particularly in the context of viral genomics. Despite limitations inherent to the simplicity of its main search algorithm, its preprocessing, specific tracking, and large genome fragmentation features, combined with an intuitive graphical interface, make it a useful resource for genomic research.

The presented case studies validate its applicability in both basic and biomedical research.

